# Studies in Particle Sorting by Paramecium Cilia Arrays

**DOI:** 10.1101/109470

**Authors:** Richard Mayne, James Whiting, Gabrielle Wheway, Chris Melhuish, Andrew Adamatzky

## Abstract

Motile cilia are cell-surface organelles whose purposes, in ciliated protists and certain ciliated vertebrate epithelia, include generating fluid flow, chemosensation, mechanosensation and substance uptake. Certain properties of cilia arrays, such as beating synchronisation and manipulation of external proximate particulate matter, are considered emergent, but remain incompletely characterised despite these phenomena having being the subject of extensive modelling. This study constitutes a laboratory experimental characterisation of one of the emergent properties of motile cilia; microparticle manipulation. The work demonstrates through automated videomicrographic particle tracking that interactions between microparticles and somatic cilia arrays of the ciliated model organism *Paramecium caudatum* constitute a form of rudimentary ‘sorting’. Small particles are drawn into the organism’s proximity by cilia-induced fluid currents at all times, whereas larger particles may be held immobile at a distance from the cell margin when the cell generates characteristic feeding currents in the surrounding media. These findings can contribute to the design and fabrication of biomimetic cilia, with potential applications to the study of ciliopathies.

## 1 Introduction

Cilia are hair-like cell-surface organelles whose functions, in multi-ciliated cells, include mechanosensing, chemosensing, substance uptake and establishment of fluid flow. The latter is achieved by a rhythmic, whip-like beating motion that serves to create local fluid currents; in single-celled ciliated organisms such as *Paramecium caudatum* (Fig. 1), this mechanism contributes to the generation of motile force. Multiciliated epithelia in vertebrates have a number of important functions in development and homeostasis. Motile cilia on the primitive node of developing amniotes establish leftward fluid flow to establish the left-right body axis during development. Defects in motile nodal cilia lead to development of laterality defects, including heterotaxia, dextrocardia, isomerism and situs inversus. Situs defects are sometimes accompanied by chronic respiratory infections in patients with Kartagener Syndrome, caused by defects in motile cilia which line the respiratory tracts and reproductive tracts and maintain mucous movement, and normal function in these systems. Motile cilia also line the ependymal cells of the ventricles of the brain to maintain flow of cerebrospinal fluid, and when these cilia are defective, severe conditions such as hydrocephaly can result. Collectively, the human conditions associated with defects in motile cilia are termed the motile ciliopathies, and contribute to a significant health burden. Whilst the incidence of individual ciliopathies suggests these conditions are rare (primary ciliary dyskinesia, which is caused by lack of motility in motile cilia has an incidence of 1 in 10,000 live births), defects in motile cilia are likely to be common causes of respiratory problems, infertility and hydrocephalus, with a considerable health and financial burden [1].

**Fig 1.**
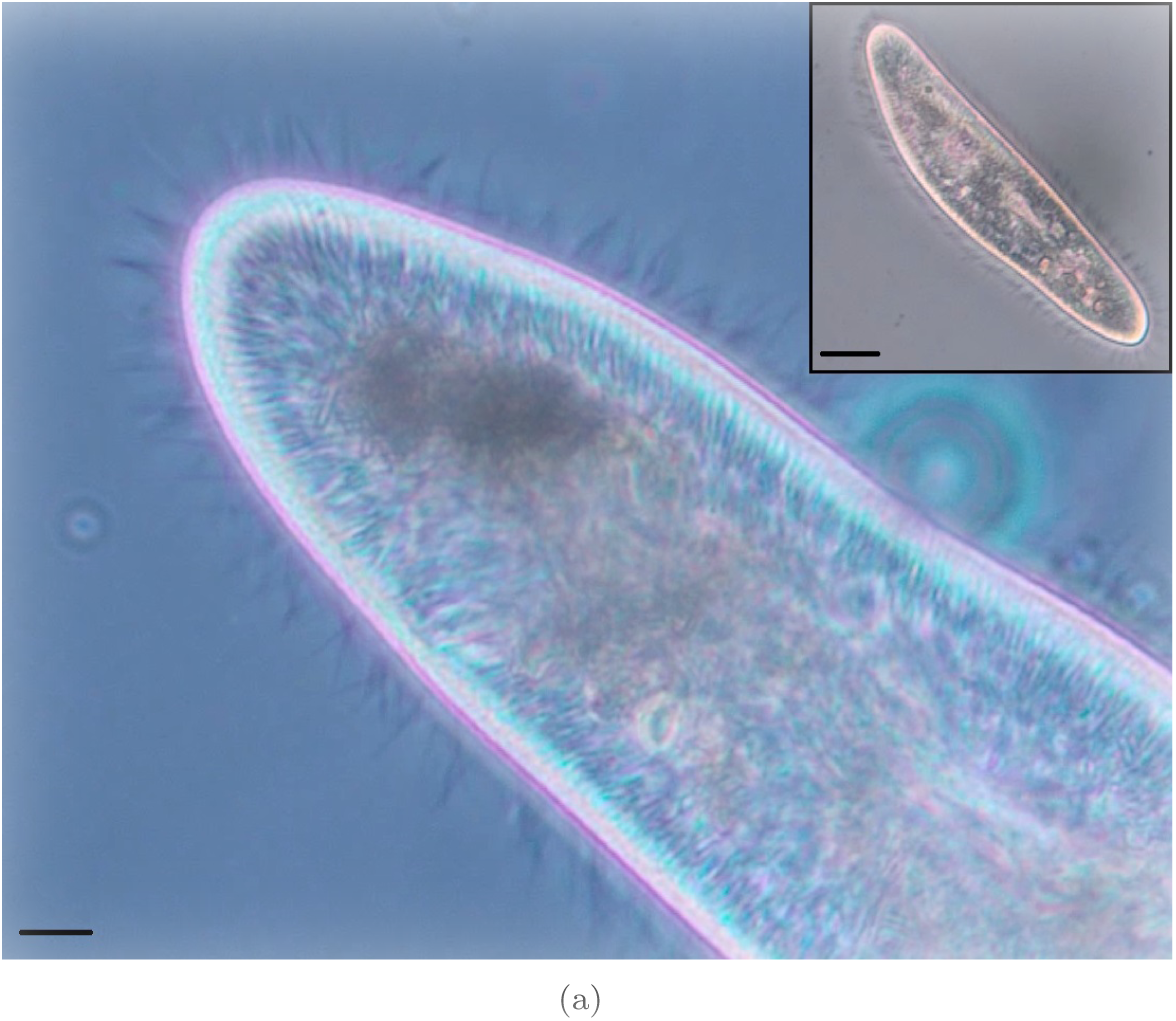
Phase contrast photomicrographs of the *P. caudatum* cell. (Main) Anterior end, in which cilia (spinulose protuberances) may be observed coating the organism. Scale bar 10 *μ*m. (Inset) Entire cell, scale bar 25 *μ*m.

Despite ciliary ultrastructure being relatively well-characterised, there is currently still some doubt concerning how emergent, synchronous processes arise from the collective actions of individual cilia [2], which thus impedes efforts towards developing treatments for ciliopathic diseases. Furthermore, certain characteristics of ciliary beating, such as their lack of a well-defined or centralised controller and their ability to collectively establish asynchronous travelling (metachronal) waves, have attracted the attentions of biologists and engineers alike as prime examples of emergent behaviour. Biomimetics research in the field of ciliary dynamics is desirable from several perspectives, as end-point outcomes include developing a deeper understanding of human disease processes, generation of artificial cilia/*in silico* biomedical research platforms and delineation of principles for bio-inspired ‘smart actuator’ hardware.

The purpose of this investigation is to experimentally characterise an emergent behaviour, particle manipulation, exhibited by the ciliated protistic model organism *P. caudatum*, with the aim of informing the design of biomimetic artificial cilia and bio-inspired actuators.

Several computational models of ciliary dynamics have attempted to describe various emergent ciliary processes [3–5] and substantial efforts have already been expended towards fabricating biomimetic cilia. Functional prototypes of artificial cilia include: individual macro-scale cilia that bend by virtue of being composed of an electro-active polymer [6]; self-assembling superparamagnetic bead chains that may be induced to beat through applying an external magnetic field [7]; arrays of vibrating motor-driven macro-scale silicon cilia [4]; cilia-inspired microelectromechanical systems devices [8]; cilia-like micromechanical electrostatic actuators for use within microfluidic circuitry [9]; decentralised cellular automaton controllers of cilia-inspired paddles in swimming robots [10]; photosensitive liquid crystal microactuators [11]. Whilst the biomimetic cilia developed to date are all ingenuitive examples of bio-inspired design, none have exhibited the desirable emergent features previously described and very few mimic more than a single aspect of cilia design/control.

Particulate manipulation by *P. caudatum* cilia is a means of enhancing its feeding mechanisms: the organism only feeds (i.e. ingests food particles) when it is swimming slowly or entirely static. As a suspension-feeding organism, *P. caudatum* harvests bacteria and small protozoa from the fluid medium it inhabits via a specialised phagocytotic vesicle within a buccal cavity which is connected to the organism’s environment via a mouth-like structure (cytostome) connected to an internal vestibule, both of which are recessed within an oral groove (peristome). Collectively, the oral groove and vestibule form the cytopharynx [12, 13]. The peristome has its own ciliary field which is denser and beats at a higher frequency than the organism’s somatic cilia. These peristomal cilia are known to draw particulate-laden fluid downwards into the cytopharynx where parallel arrays of cytostomal ciliature (quadrulus and dorsal and ventral peniculi) separate and concentrate the particulate matter prior to its ingestion, whilst ejecting fluid and particles which do not pass into the cytostome [14]. It was once thought that the organism could selectively internalise specific particulates based on ciliary sensing of their chemical content [12, 15], but recent research suggests that the parallel arrays of cilia act in a sieve-like manner, i.e. smaller particles are mechanically filtered into the oral apparatus by a dense, overlapping ciliary field [13, 16–18].

Although these mechanisms of particle manipulation within the cytopharynx have been studied extensively [14, 16, 19, 20], several research questions remain unanswered pertaining to the manner in which particulate matter is manipulated extra-cytopharyngeally. These include (but are not limited to) what contribution (if any) do the somatic cilia make to particle manipulation; are differently-sized objects all manipulated in the same manner and; how does particle manipulation differ when the organism is feeding and not feeding?

Answering these questions is vital to improving understanding of the process of particle manipulation which is necessary to facilitate its experimental manipulation/hardware emulation. The novelty of this investigation lies within examining the behaviour rather than its underlying mechanisms, as emergent behaviour is significantly more difficult to describe in terms of the individual interactions which collectively generate them. Presented here is a microscopical study which focuses on observing the dynamics of particulates in the presence of *P. caudatum* cells.

## 2 **Materials and Methods**

### 2.1 Microorganism Culture

*Paramecium caudatum* were cultivated at room temperature (21–24°C) in Chalkley’s medium enriched with 10 g of desiccated alfalfa and 25 grains of wheat per litre. Culture vessels were exposed to a day/night light cycle but were not kept in direct sunlight. Organisms used in experiments were harvested in log growth phase.

### 2.2 Organism Immobilisation

Organisms were immobilised using a method designed to preserve normal cilia dynamics as much as possible. Approximately 1000 organisms were isolated via centrifuging at 35 G for 5 minutes before being transferred to 10 ml of fresh medium. 50 *μ*l of 40 mg/ml starch-coated, 200 nm diameter magnetite (iron II/III oxide) nanoparticles (Chemicell GmBH, Germany) were then added to the medium and the organisms were incubated for 24 h. The organisms were found to consume the nanoparticles (amounting to volumes of up to 25 % total cell volume) which rendered them paramagnetic; this allowed the organisms to be immobilised for microscopical study on well slides by placing a 40 × 25 mm, 1.3 T neodymium magnet in close proximity (Fig. 2).

**Fig 2.**
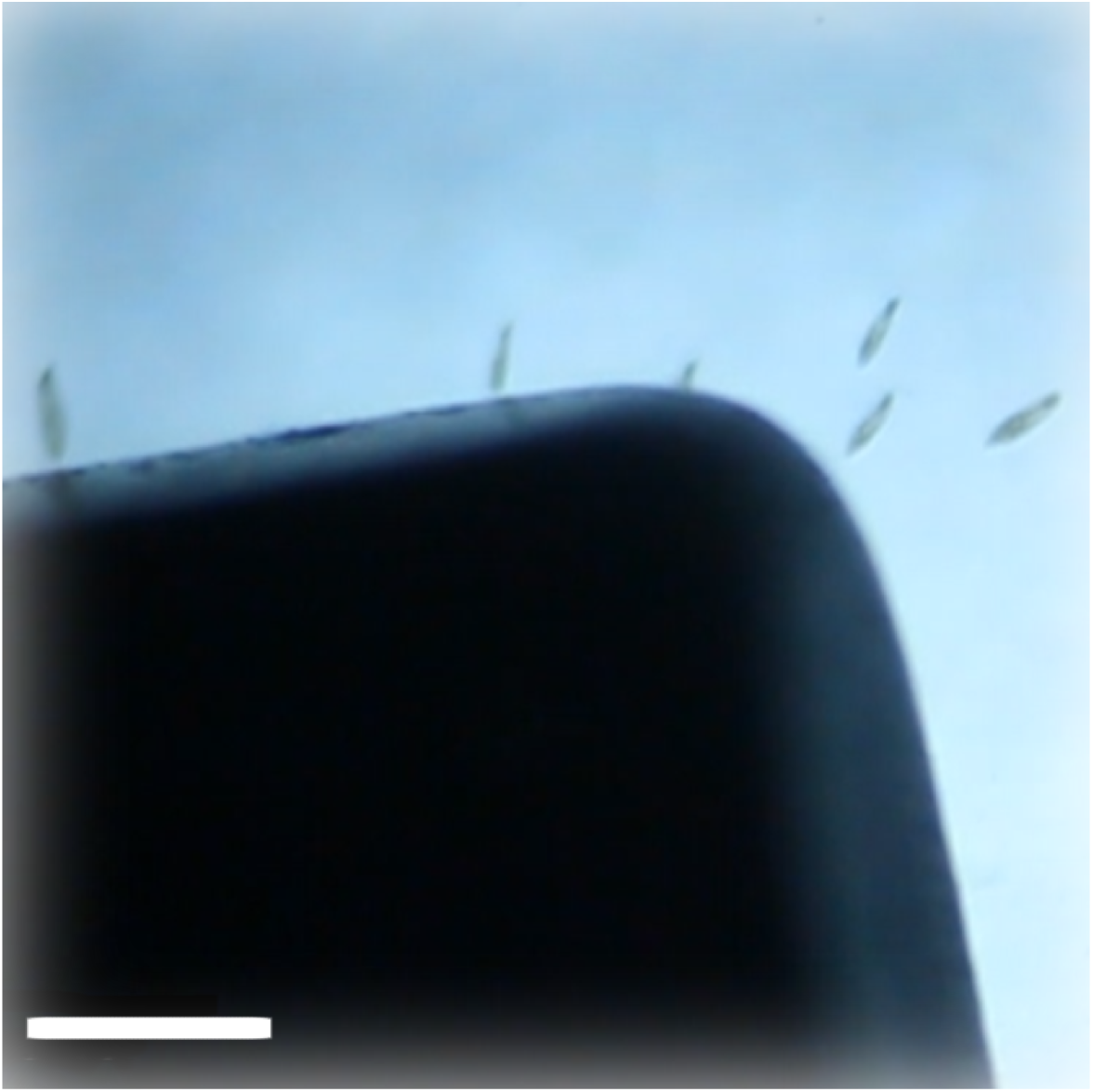
Photomicrograph showing magnetite nanoparticle-treated *P. caudatum* being drawn to a magnet. Scale bar = 500 *μ*m.

By varying the distance between the magnet and the well, the organisms could be held in either sessile (feeding; magnet further away) or attempting to migrate (not feeding; magnet closer) states; these phases were distinguished by the similarity of the ciliary beating-induced fluid currents around the organism to those of static/slow moving and rapidly moving organisms in control cultures.

### 2.3 Particle Manipulation

In order to observe ciliary manipulation of dispersed particulates, single organisms were transferred to wells containing fluorescent carboxylated latex microparticles (Fluospheres, Thermo Fisher, USA). The following varieties of particle were used:

1. 0.2 *μ*m diameter (505/515 nm excitation/emission), 2 × 10^6^ particles per ml.
2. 2.0 *μ*m diameter (580/605 nm), 2 × 10^6^ particles per ml.
3. 15 *μ*m diameter (580/605 nm), 1 × 10^6^ particles per ml.

The choice of sizes was informed by Ref. [16], which states that ca. 2 *μ*m is the optimal size of particle for *P. caudatum* with regards to the rate at which they may be internalised; the other two varieties were chosen for being (approximately) an order of magnitude bigger/smaller in size. All latex spheres were pelleted as per manufacturer instructions and resuspended in deionised water in order to remove preservatives. The organisms were allowed several minutes to equilibrate to their new environment before being observed. Observations were made with a Zeiss Axiovert 200M inverted microscope; concurrent brightfield and fluorescence observations were made using a Perkin Elmer UltraView ERS FRET-H spinning disk confocal microscopy system. Optical section depth was maintained at 0.3 *μ*m in order to minimise the visualisation of particle movements outside of the central field of view, including particle movements within the oral cavity. Video footage was post-processed with Volocity 5.3 in which colour assignment and brightness/contrast adjustments were made.

### 2.4 Video analysis

Video analysis was performed using a custom Matlab (Mathworks, USA) script. Videos were first cropped to an area of approximately 300 *μ*m^2^. RGB images were imported from the video frame-by-frame for sequential analysis and particle positioning. For each particle video set, the respective colour was isolated from the RGB image, whereupon the data for each frame was converted to a JPEG image file for further analysis. To detect the position of each particle, a Laplace template of a Gaussian filter was defined before being convolved over the image; the size of the filter was iteratively determined by visual feedback of the user for optimal particle detection for each of the 3 particle sizes. Each image in a video sequence was subject to filtering. After particle detection on every frame had occurred, the positional data was passed to a bespoke Kalman filter which accurately estimates the position of the particle across each frame using the data from the full time-series of particle positions to predict and confirm the movement of each particle. From this it is possible to measure the speed of each particle in a noisy video, creating a dataset of particle speed and position for each particle size.

Each video was calibrated using a haemocytometer slide, from the fixed grid size, a pixel to micrometre ratio was determined, providing accurate analysis of particle speed. While the script ran, frame-by-frame images were shown on screen allowing visual validation of positional tracking by the authors; erroneous particle tracking was removed after convolution but before analysis was performed. Average speed was calculated for each particle passing through the fluidic vortex around the organism.

## Results

Two distinct patterns of particle movement around *P. caudatum* cells were identified which corresponded to two modes of fluid current (Figs. 3 and 4, S1–3). One pattern corresponded to feeding behaviour (Fig. 3), as indicated by the organism requiring only mild magnetic restraint and the similarity of the appearance of the fluid currents with control organisms, whereas the other appeared to be associated with migration due to the organisms exhibiting this behaviour requiring greater strength of magnetic restraint (Fig. 4).

**Fig 3.**
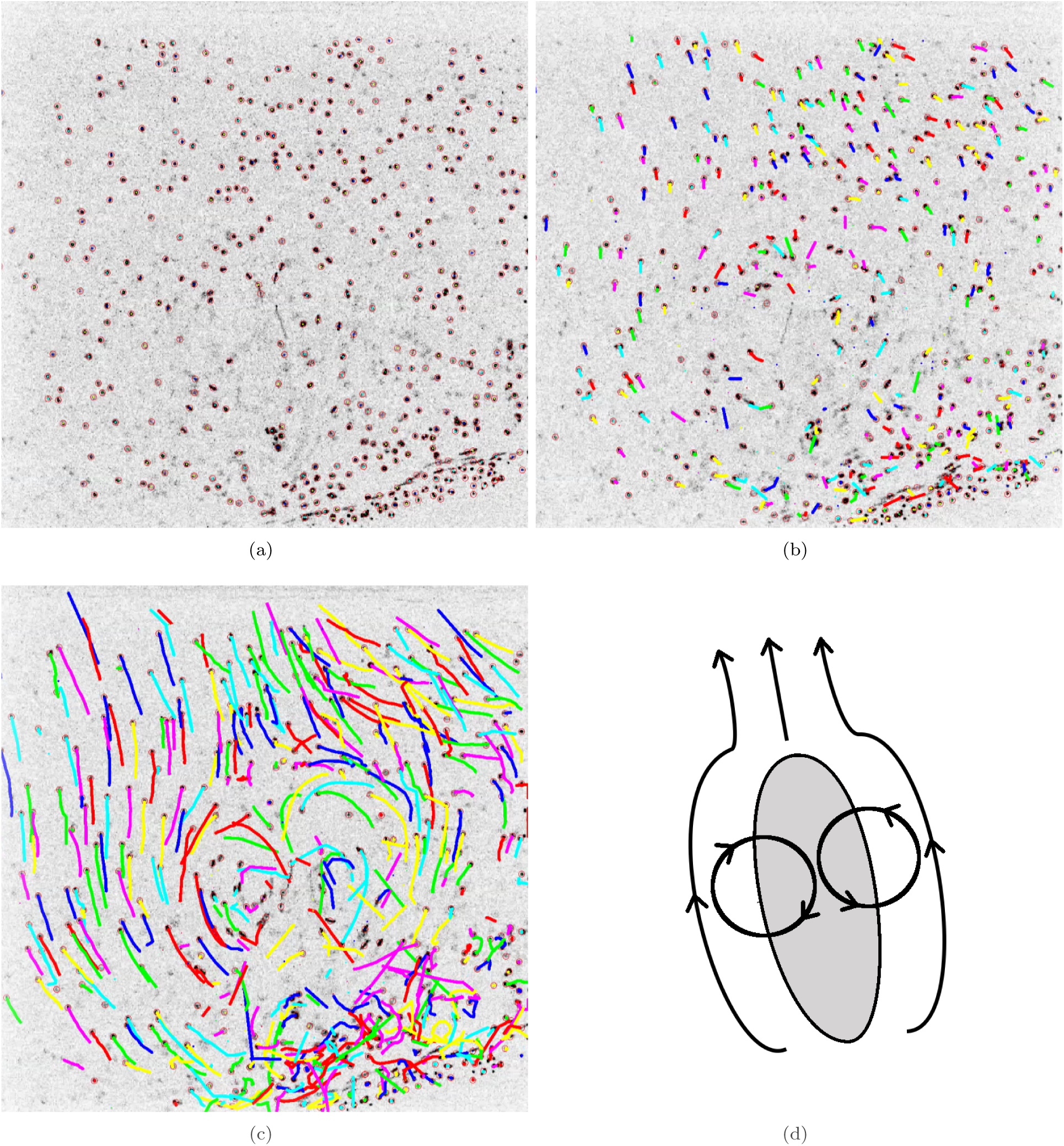
Particle tracking from video footage of a feeding, magnetically-restrained *P. caudatum* cell in the presence of 2 *μ*m fluorescent particles. (a–c) Three frames of output from particle tracking script. Red circles indicate a detected particle and movement from their location in the previous frames is highlighted with variously-coloured contrails. (d) Diagram derived from 〈*c*〉 showing cell’s position (grey oblong) and the direction of particle movements (arrowed lines).

**Fig 4.**
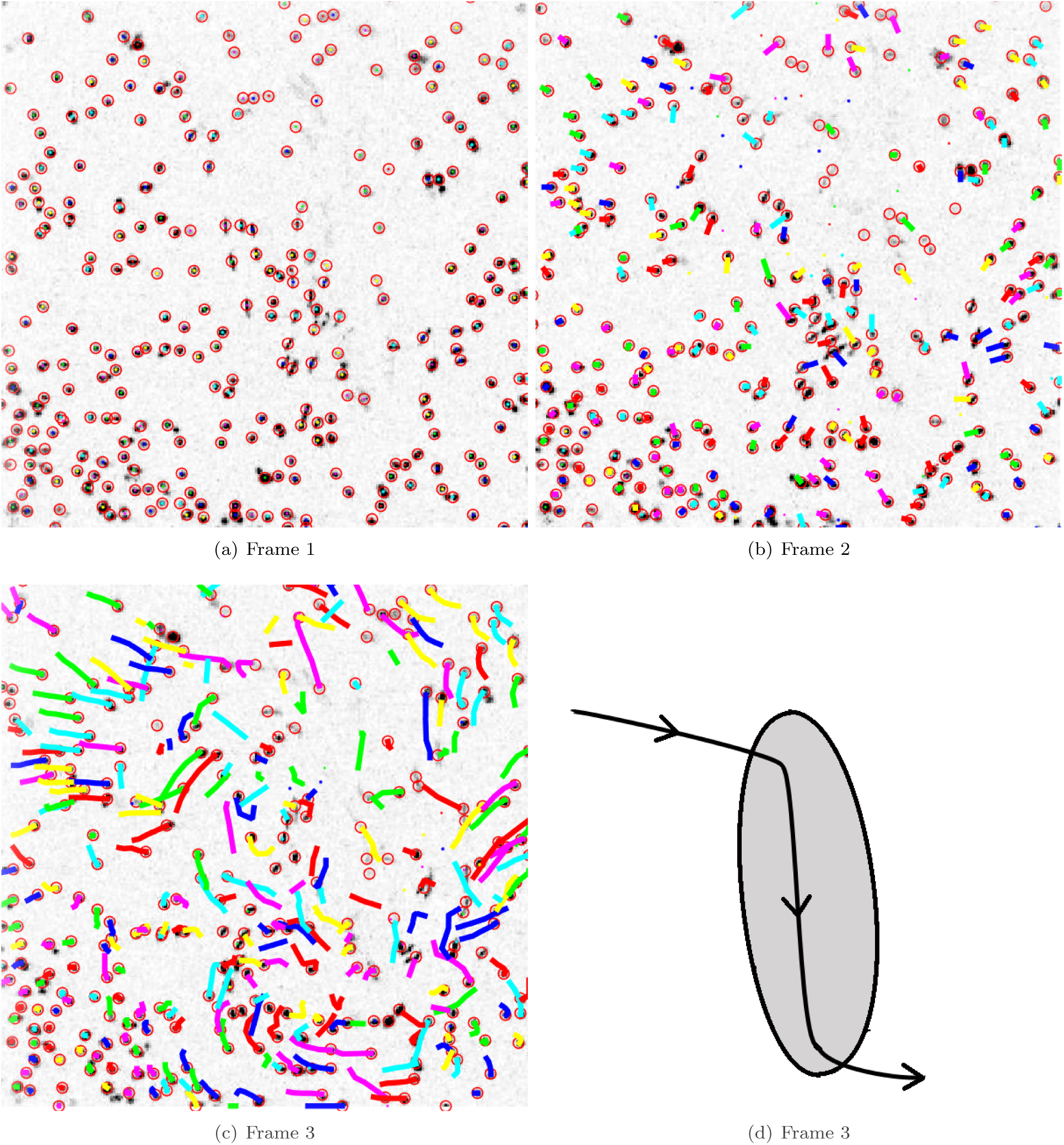
Particle tracking from video footage of an actively migrating, magnetically-restrained *P. caudatum* cell in the presence of 2 *μ*m fluorescent particles. (a–c) Three frames of output from particle tracking script. Red circles indicate a detected particle and movement from their location in the previous frames is highlighted with variously-coloured contrails. (d) Diagram derived from 〈*c*〉 showing cell’s position (grey oblong) and the direction of particle movements (arrowed lines).

Both 0.2 *μ*m and 2.0 *μ*m particles were observed to be drawn into close proximity with paramecia in both feeding and motile states. The average speeds of transport for each particle variety in organisms exhibiting feeding behaviour are shown in Table 1 (full data and statistics are included in S4). Mean particle speeds for each variety were significantly different from abiotic controls, i.e. separate from Brownian motion. Mean transport speeds for 0.2 and 2.0 *μ*m particles were significantly different to 15 *μ*m particles but were not significantly different from each other. The effective distance at which *P. caudatum* cells could draw 0.2 *μ*m and 2.0 *μ*m particles in typically exceeded the range of the visualisation range (300 *μ*m^2^), although the speed at which they moved within this range was not uniform.

**Table 1.** Data to show the average speed of particles within the immediate vicinity of *P. caudatum* cells in both feeding and migrating states.

| Particle Size | Mean particle speed ($\mu\text{m s}^{-1}$) | Control ($\mu\text{m s}^{-1}$) |
| --- | --- | --- |
| 0.2 $\mu\text{m}$ $\bar{x}$ | 137.89 | 28.53 |
| $\sigma$ | 44.50 | 12.81 |
| 2.0 $\mu\text{m}$ $\bar{x}$ | 137.97 | 21.03 |
| $\sigma$ | 27.16 | 4.57 |
| 15 $\mu\text{m}$ $\bar{x}$ | 175.21 | 7.76 |
| $\sigma$ | 56.75 | 2.52 |

15 *μ*m particles were not captured by the fluid vortices created by actively migrating paramecia, as was indicated by their movements not being detected by the tracking script. When the cells happened to swim directly into 15 *μ*m particles, they were only displaced by the passage of the organism rather than being drawn into its vortices and consequently ejected in a fluid contrail. Larger particles were, however, drawn into the fluid vortices generated by feeding behaviour.

Following being drawn into the organisms’ feeding currents, 15 *μ*m particles were regularly, but not invariably, observed to become static approximately 10 *μ*m from the cell margin (Fig. 5). The duration of this effect was highly variable, ranging from a few seconds to several minutes. No means of physical connection between particles and cells were observed. Fluid vortices were still visible surrounding organisms exhibiting this behaviour. Eventually, the immobilised particles were released, entered the fluid vortex and were propelled away from the cell.

**Fig 5.**
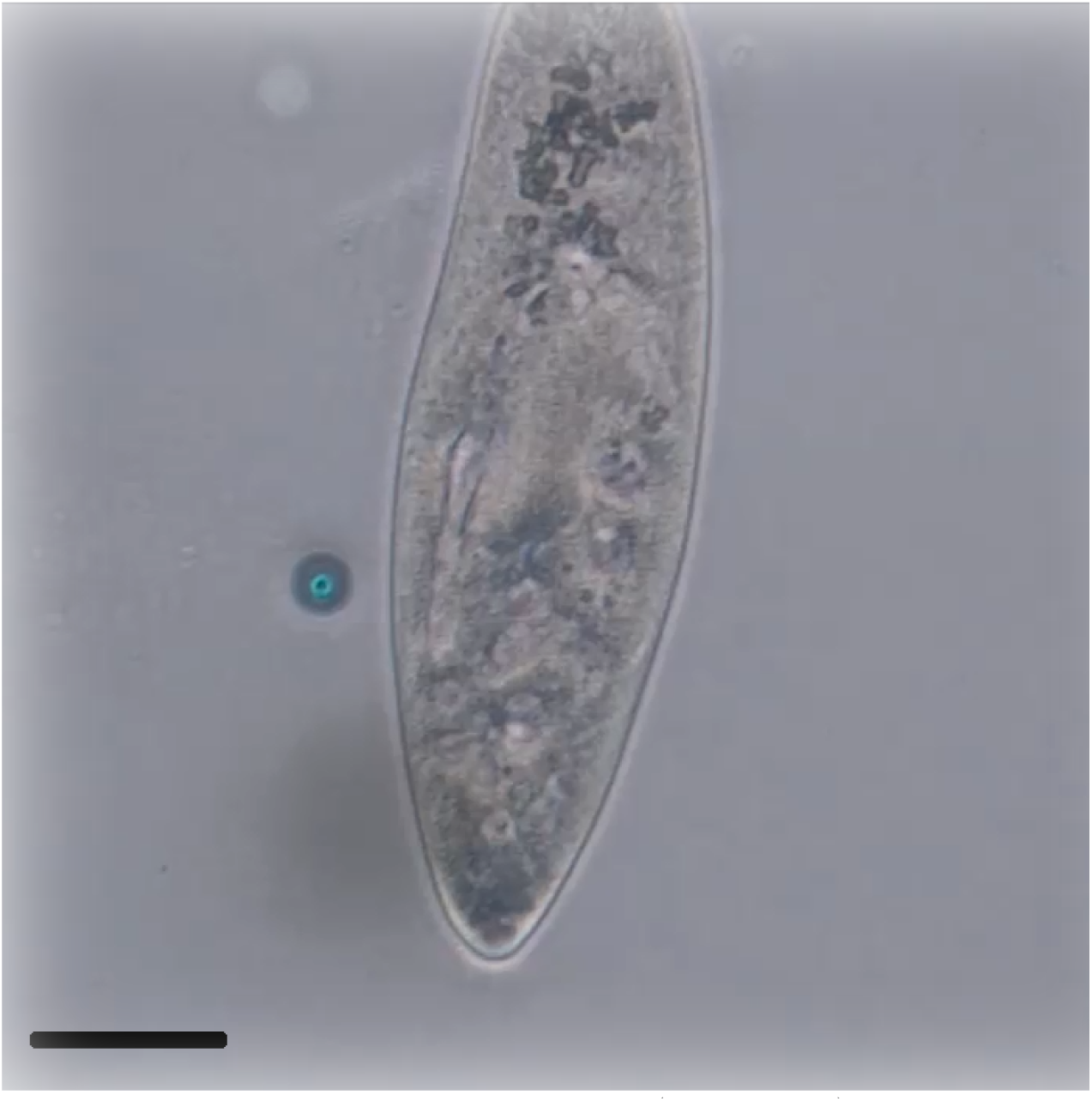
Phase contrast micrograph showing a 15 *μ*m particle (blue object) suspended in close proximity with a *P. caudatum* cell that was generating a ‘feeding’ pattern fluid vortex (S3). No physical method of attachment (such as a trichocyst) is easily visible between cell and particle. Scale bar = 50 *μ*m.

Both 0.2 *μ*m and 2.0 *μ*m particles were observed within the organisms’ intracellular vesicles (visualised via fluorescence), indicating that they had been consumed; no intracellular 15 *μ*m particles were observed.

## 4 Discussion

The results presented indicate that the activities of *P. caudatum* somatic cilia result in the emergence of discrimination between the particles that are drawn into their proximity: large particles may become suspended approximately one cilium’s length away from the surface of the cell (10 *μ*m [21]), whereas smaller particles are drawn in, presumably towards the oral apparatus for internalisation. With regards to the theme of this article, this behaviour can be interpreted as differential manipulation of substances according to some intrinsic property of the material being sorted or, more concisely, ‘sorting’. This represents a key finding of this study as *P. caudatum* cilia has only been described as being capable of mechanical, sieve-like sorting within the oral cavity. Presented here, conversely, is evidence of a further form of discrimination between particulates which is coordinated by the somatic cilia.

The mechanisms behind the anchorage of latex particles to the surface of the cell are unclear from our results. We propose two possible methods: firstly, it is possible that the process is mediated by trichocysts. Trichocysts are needle-like secretory organelles whose function is still partially unelucidated, but are considered to be primarily (if not exclusively) a defence mechanism [22]. Although no trichocysts were observed interacting with adjacent particles, it is possible that the microscopical techniques used (phase contrast/fluorescence at ×100 magnification) were not adequately suited to the visualisation of individual trichocysts. Secondly, large particle immobilisation could be due to the formation of a double layer as a function of electrostatic interactions between the cell membrane, medium components and the latex particles.

In either case, this mechanism appears to enhance foraging behaviour as fluid currents continue unabated when 15 *μ*m particles are suspended, which presumably facilitates stripping smaller particulates (bacteria, small protists) from immobilised detritus. Indeed, *Paramecium* spp. have been previously described as being capable of attaching themselves to solid surfaces in order to enhance feeding [23]. Further investigation of this phenomenon is an important area for further study as characterisation of its underlying mechanisms are necessary to inform the design of biomimetic cilia. If attachment to large particles is mediated by trichocysts, this implies the involvement of a sensing loop, whereas the electrostatic hypothesis requires no sensing but would instead require careful engineering of cilia surface properties.

Regardless, it is clear that the intrinsic properties of *P. caudatum* cilia participate in manipulating particulate material. Lacking a centralised controller, a significant portion of the sorting task is outsourced to the ciliary morphology; for example, the characteristics of feeding currents are a direct function of ciliary morphology, compliance and orientation. This observation is complementary to recent advances in morphological computation and entity embodiment [24]; hence, we emphasise that artificial cilia designs should focus upon these principles. Whilst recent advances in biomimetic cilia have begun to make use of soft robotics approaches [4, 6], none have yet capitalised upon anything but a compliant body.

Hardware-based comparisons of sorting by cilia possessing or lacking sensing abilities would further contribute to our understanding of the biological basis of this phenomenon. It is still unclear how these processes are synchronised in the absence of a centralised controller. Our previous work has indicated that sensorimotor coupling is a function of a cell’s cytoskeleton [25] and we therefore recommend that this is a logical approach to further research in the field.

Fenchel’s extensive observations on the optimum size of particles to be ingested by the oral apparatus of various ciliates [16] indicated that particles smaller than ≤ 0.2 *μ*m are sufficiently small to fall between the interciliary gaps and are hence ejected in the organism’s fluid contrails rather than being passively filtered towards the oral aperture. Our observations of 0.2 *μ*m particles having being internalised apparently contradict this, although no measures were taken to prevent the latex spheres from aggregating or otherwise binding with other particulates in the culture medium. Methods of particle sorting by *P. caudatum* cilia are presented algorithmically in Fig. 6.

**Fig 6.**
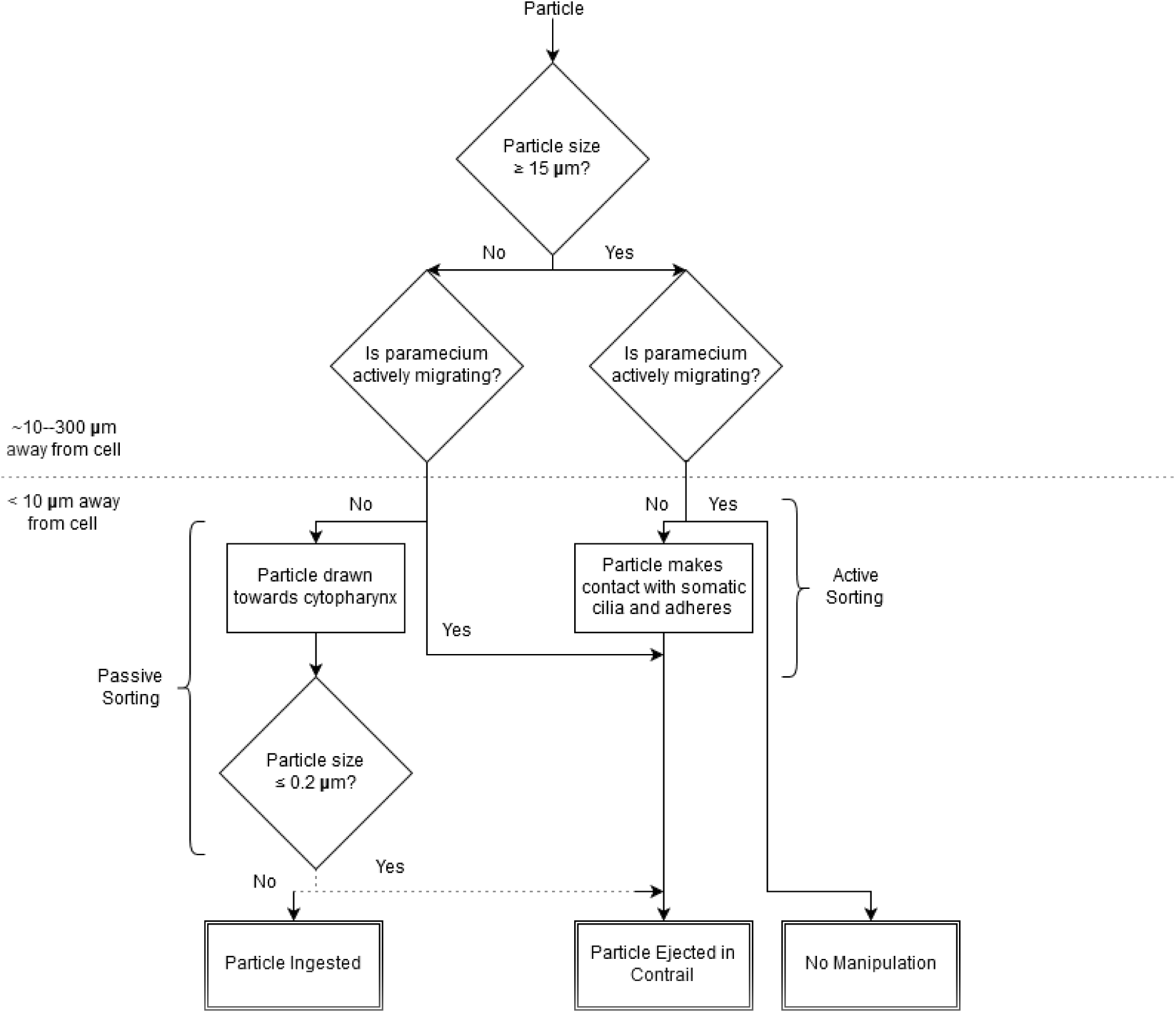
Algorithm to show process of particle sorting by *P. caudatum* ciliature and cilia-derived fluid movements. Dashed arrows represent data indicated in Ref. [16] but not observed in this study.

The observation of two distinct patterns of fluid movement about the *P. caudatum* cell is of note here as an exhaustive literature search only identified the documentation of the ‘feeding’ pattern shown in Fig. 3 [26], despite the two patterns being easily discernible when observing cultures via phase contrast microscopy. The implications of the organism’s ability to exhibit two varieties of fluid current are, although unsurprising given the physiological states they represent, important observations for those involved in the design of biomimetic micro- and nano-machines, considering that cilia-like nano-motors have recently been described as a feasible technology [27].

This work contributes to our understanding of one of the emergent properties of motile cilia; particle sorting. A thorough understanding of these properties of cilia arrays is important for the design of biomimetic cilia, which have potential applications in industry, for particle sorting in liquid suspensions, and for development of prosthetic cilia as a treatment for immotile ciliopathies, such as primary ciliary dyskinesia.

## Acknowledgements

### 5 Appendices

#### 5.1 Acknowledgements

The authors extend their sincerest thanks to Dr. David Patton for his various critical insights.

## 5.2 Supplementary Information Captions

- **S1 Video:** Confocal videomigrograph of demonstrating the movement of 2.0 *μ*m particles about a *P. caudatum* cell, corresponding to the tracking output in Fig. 3.
- **S2 Video:** Video output from stills shown in Video 1/Fig. 3.
- **S3 Video:** Brightfield videomicrograph demonstrating the movement of a 15 *μ*m particle about a *P. caudatum* cell.
- **S4 Spreadsheet:** Measurements and statistical data for particle speed tracking.

